# Correcting MEG artifacts caused by overt speech

**DOI:** 10.1101/2020.11.28.402008

**Authors:** Omid Abbasi, Nadine Steingräber, Joachim Gross

**Author notes:** **Corresponding author:** Omid Abbasi, Institute for Biomagnetism and Biosignalanalysis, University of Münster, Malmedyweg 15, 48149 Münster, Germany.

## Abstract

Recording brain activity during speech production using magnetoencephalography (MEG) can help us to understand the dynamics of speech production. However, these measurements are challenging due to the induced artifacts coming from several sources such as facial muscle activity, lower jaw and head movements. Here, we aimed to characterise speech-related artifacts and subsequently present an approach to remove these artifacts from MEG data. We recorded MEG from 11 healthy participants while they pronounced various syllables in different loudness. Head positions/orientations were extracted during speech production to investigate its role in MEG distortions. Finally, we present an artifact rejection approach using the combination of regression analysis and signal space projection (SSP) in order to correct the induced artifact from MEG data. Our results show that louder speech leads to stronger head movements and stronger MEG distortions. Our proposed artifact rejection approach could successfully remove the speech-related artifact and retrieve the underlying neurophysiological signals. As the presented artifact rejection approach was shown to remove artifacts induced by overt speech in the MEG, it will facilitate research addressing the neural basis of speech production with MEG.

## 1. Introduction

Our everyday life and especially social interactions are greatly affected by verbal communication and, consequently, the question of how our brain is able to produce and understand spoken language has been the focus of many studies in the field of cognitive neuroimaging. fMRI studies have the spatial resolution to potentially provide a spatially detailed account of brain areas involved in speech perception and production (Hickok, 2012; Hickok and Poeppel, 2016). However, scanner noise and sensitivity to head movements caused by overt speech make fMRI less than ideal – a limitation that is further aggravated by the limited temporal resolution of fMRI. Instead, magnetoencephalography (MEG) and electroencephalography (EEG) provide excellent temporal resolution and are completely silent recording techniques (Gross, 2019). This makes them ideal candidates to non-invasively study the dynamics of speech perception and production. However, particularly speech production studies are challenging due to the induced artifacts coming from several sources such as facial muscle activity, lip and eye movements, and head movements (Ewald et al., 2012). Therefore, earlier studies have avoided recording neurophysiological activities during speech production. Some tried to simply circumvent the induced artifacts by delaying continuous speech, using silent naming, or manual responses (Eulitz et al., 2000; Ewald et al., 2012; Schmitt et al., 2000). However, these approaches compromise our ability to directly investigate neural activity underlying speech production.

Recently, some studies have used MEG and EEG to study speech production (Alexandrou et al., 2016; Bourguignon et al., 2020; De Vos et al., 2010; Ewald et al., 2012; Ganushchak et al., 2011; Llorens et al., 2011; Masaki et al., 2001; Meyer, 2018; Munding et al., 2016). While EEG is less affected by head movement compared to MEG, source localisation is generally thought to be more precise with MEG (Gross, 2019).

Therefore, here we focus on the use of MEG for studying speech production. While artifacts are an obvious problem for these studies, they have so far not been carefully described. In this study, we seek to comprehensively characterise speech artifacts in MEG and evaluate the performance of correction techniques with the aim to optimise source localisation.

Several artifact correction approaches have been used to reduce speech artifacts in MEG signals. Temporal signal space separation (tSSS) uses expansions of spherical harmonic functions to separate MEG signal components originating inside the sensor array from those originating elsewhere (Taulu and Simola, 2006). While this is not suitable to remove facial muscle artifacts (which originate inside the sensor array), tSSS can be used to compensate for head movements and suppress external artifacts. However, tSSS was developed for a specific type of MEG system (Elekta) and is not available for other MEG systems.

In addition, independent component analysis (ICA) has been used to remove speech-related artifacts from the recorded MEG data (Alexandrou et al., 2017; Bourguignon et al., 2020; Laaksonen et al., 2012; Liljeström et al., 2015b; Ruspantini et al., 2012). These studies used ICA to decompose data from the sensor level to the component level. Then, components related to artifacts were detected and removed from data. ICA is in principle also suitable for removing muscle artifacts but requires an accurate selection of artifactual components (Gross et al., 2013).

Besides head movements and muscle artifacts, movements of the lower jaw are another prominent source of artifacts during speech production. Most participants have tooth fillings, retainers, or other dental work that often has residual magnetic activity even after demagnetization. This can lead to strong artifacts that are independent of head movements. Since head movements are tracked by coils attached to the scalp, lower jaw movements are not corrected by head movement correction methods.

Here, we present an analysis pipeline that aims to correct these different types of artifacts using non-commercial methods that are applicable to data from any MEG system. We start by characterising the dynamics of head movement and recorded facial EMG activity relative to the produced speech envelope and the corresponding artifacts in the MEG signal. Finally, we present and evaluate a system-independent approach to remove the artifacts. This approach uses regression analysis for head movement correction and signal space projection (SSP) for remaining artifacts such as those arising from lower jaw movements.

To test this approach, we recorded MEG data from eleven participants who were instructed to pronounce different syllables in various loudnesses. We demonstrate that the head movement level is directly linked to the loudness level and leads to MEG distortion. Our proposed artifact rejection scheme was able to remove the movement-related artifact from MEG and retrieve neurophysiological signals.

## 2. Method

In this study, we analysed MEG data from two different experiments *(EXP1* and *EXP2). EXP1* was conducted in order to characterise movement-related artifacts induced during speech production and further to investigate the feasibility of removing these artifacts from the recorded MEG data. Moreover, we also analysed data from a larger experiment *(EXP2)* in order to evaluate if our artifact rejection approach removes neurophysiological information.

### 2.1 Participants

We recruited eleven healthy volunteers (6 males, mean age 25.4 +/- 2.7 y, median 25, range [22 31]) for *EXP1* and 25 healthy volunteers (13 males, mean age 24.9 +/-2.8 y, median 25, range [21 32]) for *EXP2* from a local participant pool. The study was approved by the local ethics committee (University of Münster) and conducted in accordance with the Declaration of Helsinki. Prior written informed consent was obtained before the measurement and participants received monetary compensation after the experiment.

### 2.2. Recording

MEG, electromyogram (EMG), and speech signals were recorded simultaneously. A 275 whole-head sensor system (OMEGA 275, VSM Medtech Ltd., Vancouver, Canada) was used for all of the recordings with a sampling frequency of 1200 Hz, except the speech recording which had a sampling rate of 44,1 kHz. Audio data were captured with a microphone, which was placed at a distance of 155 cm from the participants’ mouth, in order not to cause any artifacts through the microphone itself. Three pairs of EMG surface electrodes were placed after tactile inspection to find the correct location to capture muscle activity from the *m. genioglossus, m. orbicularis oris* and *m. zygomaticus major.* One pair of electrodes was used for each muscle with about 1cm between electrodes. A low-pass online filter with a 300Hz cut-off was applied to the recorded MEG and EMG data.

### 2.3. Paradigm

For both experiments participants were asked to sit relaxed while performing the given tasks and to keep their eyes focused on a white fixation cross. In *EXP1* they were instructed to pronounce three different syllables (‘La’, ‘Du’, ‘Pa’) and each of them at three different volume levels: normal, loud, and without sound, meaning they should only perform the mouth movement. This makes nine conditions in total. 50 trials were collected for each of them, consisting of 4s in which participants pronounced the syllable separately 3 to 4 times and 2s of rest in between trials. A color change from white to blue of the fixation cross indicated the beginning of the 4s in which participants should speak. After these 4s the color changed back to white, indicating that participants should stop speaking. 2s later the color would again change back to blue.

In *EXP2* participants listened to syllables and overt speech, which they generated themself in a previous recording. Overt speech was obtained by asking participants to answer fourteen questions, each of them for 60 seconds. For the syllables, subjects were told to speak as they normally would, using only the given syllable (‘Du’,’La’,’Pa’) for three minutes each. As in *EXP1* a color change of the fixation cross from white to blue indicated the beginning of the time period in which participants should speak and the end was marked by a color change back to white.

### 2.4. Preprocessing and data analysis

Prior to data analysis, MEG data were visually inspected for detecting jump artifacts and bad channels. A discrete Fourier transform (DFT) filter was applied to eliminate 50 Hz line noise from the continuous MEG and EMG data. Moreover, EMG data was highpass-filtered at 20 Hz and rectified. Continuous head center position and rotation were extracted from the fiducial coils placed at anatomical landmarks (nasion, left, and right ear canals) leading to six time-series representing the instantaneous location (X, Y, Z) and orientation (Ox, Oy, Oz) of the head. These time-series are in a 3-D Cartesian coordinate system defined relative to the dewar in which X, Y, and Z are from head center to anterior, left, and superior respectively and Ox, Oy, and Oz are the head rotation around X, Y, and Z-axis respectively. The wideband amplitude envelope of the speech signal was computed using the method presented in (Chandrasekaran et al., 2009).

Nine logarithmically spaced frequency bands between 100-10000 Hz were constructed by bandpass filtering (third-order, Butterworth filters). Then, we computed the amplitude envelope for each frequency band as the absolute value of the Hilbert transform and downsampled them to 1200 Hz. Finally, we averaged them across bands and used the computed wideband amplitude envelope for all the further analysis. We used speech envelope signals to extract syllable pronunciation onsets for further trial selection steps. First, the envelope signal was low-pass filtered at 5 Hz. Then, we computed the first derivative of the signal followed by z-scoring. Finally, we detected all the local peaks that have Z-value higher than 2. Our multistep preprocessing procedures are depicted in Fig. 1.

**Fig. 1.**
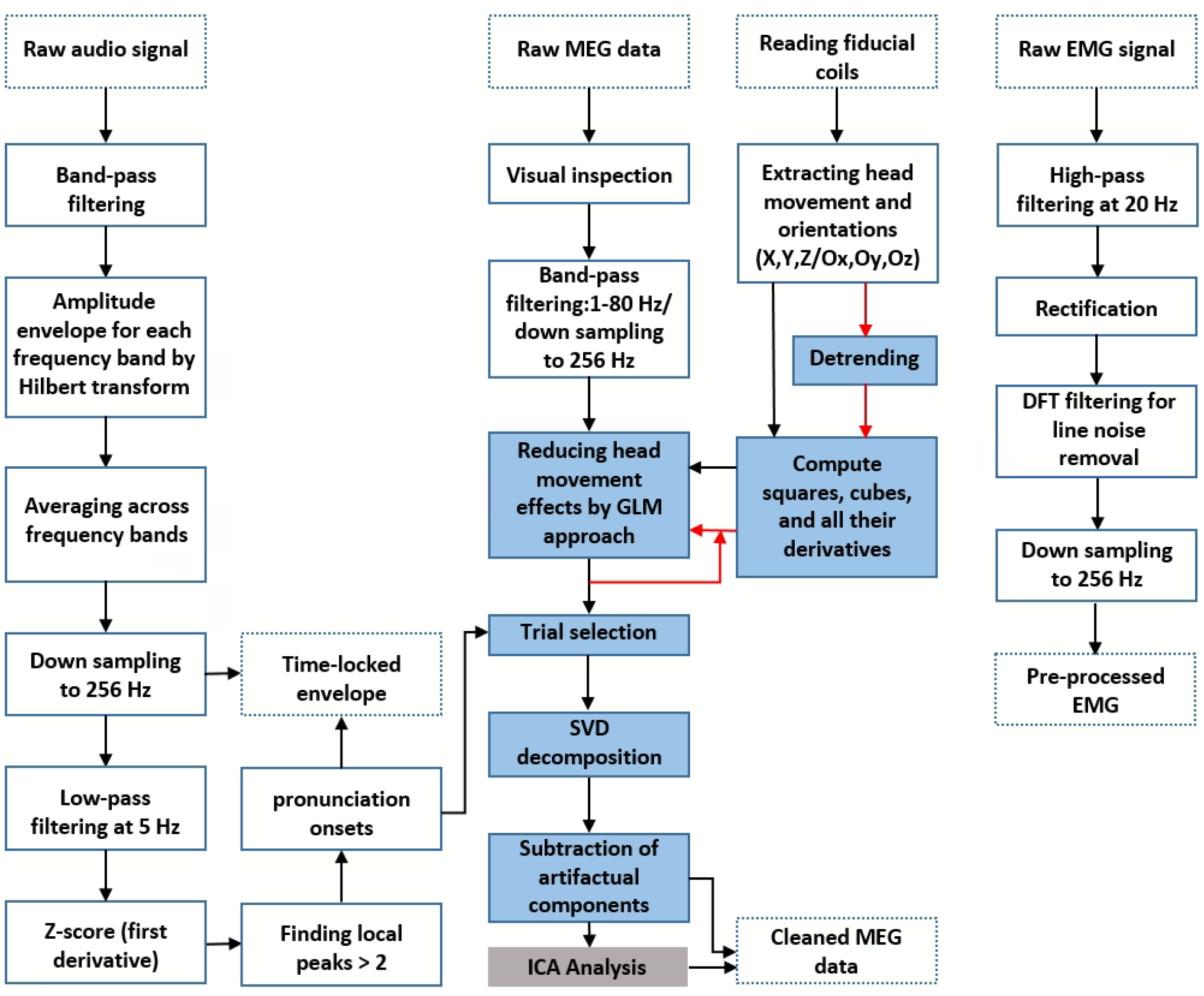
Preprocessing and artifact rejection steps. Dotted boxes indicate data input/output. Solid line boxes indicate processing steps. Blue boxes show the main steps of MEG artifact rejection. The gray box shows a complementary step for removing residual muscle artifacts according to previous studies. Red arrows indicate the second-time regression analysis after the detrending of head movement signals.

MEG, EMG, speech envelope, and head movement signals were downsampled to 256 Hz and were segmented from −1s to 4s time-locked to speech onset identified as peaks of the first derivative of the speech envelope signal. In the preprocessing and data analysis steps, custom-made scripts in Matlab R2020 (The Mathworks, Natick, MA, USA) in combination with the Matlab-based FieldTrip toolbox (Oostenveld et al., 2011) were used in accord with current MEG guidelines (Gross et al., 2013).

### 2.5. Artifact rejection

For removing the speech-related artifacts we applied the following procedure: (i) We tried to initially reduce head movement-related artifacts by incorporating the head position time-series into the general linear model (GLM) using regression analysis (Stolk et al., 2013). We considered head movement time-series, its squares, cubes, and all their derivatives in the model in order to account for both the linear and non-linear effects of head movement on the signal. We performed this step twice: once with the original head movement time-series and the second time after removing slow trends from the head-movement time-series using a third-order polynomial fit; (ii) To further remove the residual artifact, we used the SSP method implemented in the Fieldtrip toolbox (Uusitalo and Ilmoniemi, 1997). We estimated the spatial subspace containing the speech-related artifact from the MEG data using SSP. The singular value decomposition (SVD) approach was applied to the MEG data to decompose data into singular vectors (components). This method works well if it is applied to data where the artifact is maximised due to preprocessing or averaging. In our case, we averaged data time-locked to speech onset thereby enhancing the artifact in the averaged data. Therefore, the first SVD components are expected to reflect the artifact subspace. (iii) Artifactual components were detected and removed via visual inspections and mutual information (MI) analysis (Abbasi et al. 2016); (iv) finally, all remaining components were back-transformed to the sensor level. The blue boxes in Fig. 1 depict MEG artifact rejection steps.

Our conservative component selection consisted of three steps: First, visual inspection of components’ time-series: Artifactual components’ time-series were affected by movement patterns and were detectable. Second, visual inspection of the components weight distribution on the topographic maps: Components with major power distribution on temporal areas were marked as artifactual components. Third, computing mutual information between SVD components and head movement/orientation signals: MI was calculated between SVD components and each of the extracted head movement/orientation signals (X, Y, Z, OX, OY, and OZ). These MI values were subsequently averaged across all movement signals for each pair. Components with high MI values were detected as artifactual components.

### 2.6. Artifact level

In order to quantify the level of artifact and be able to compare the level of residual artifact between different artifact rejection steps, we constructed a spatial filter from the SVD of the raw data covariance matrix in *EXP1.* We only took into account the artifactual SVD components detected in our artifact rejection steps for every participant. Next, we applied the constructed filter on the data after every artifact rejection step. Finally, the artifact level was obtained from averaging the absolute values of the filtered data. Additionally, we evaluated to what extent the proposed artifact rejection approach removes neurophysiological information. For this aim, we used the spatial filters constructed from artifactual SVD components in the speech production condition to clean the artifact-free syllable perception dataset in *EXP2.* The amount of reduction in MEG components in the syllable perception condition indicates the removal of neurophysiological information since there are no speech-artifacts in the perception condition.

### 2.7. Coherence analysis

In order to quantify similarities between speech envelopes and EMG/MEG signals, we used coherence analysis. To this end, time-locked data were further segmented into short data segments (duration 2 s, overlap 0.5 s). Spectral power and cross-spectral density of these data were calculated using a multi-taper approach with a smoothing of 2 Hz (range 1 – 50 Hz). Coherence was computed for each participant between speech envelopes and EMG/MEG signals. These coherence values were subsequently averaged across all participants for each pair (EMG/MEG, Envelope) and each frequency. Finally, for each pair, the coherence values were averaged within the frequency range of 0-5 Hz.

### 2.8. Statistical analysis

In our statistical analysis, we aimed to investigate any significant effect of the speech loudness and syllables on head movement/orientation. To meet this purpose, we used linear mixed effect modeling (LMEM). We computed the averaged head movement/orientation in all directions (X, Y, Z) and orientations (Ox, Oy, Oz) for the first 4 seconds after the syllables pronunciation onset for each participant. All the average head movement/orientation values were entered into a fixed-effect model with loudness and syllable as factors. Moreover, we also investigated whether there are any significant effects of speech loudness, syllables, and EMG signal on the calculated coherence between the speech envelope and EMG signals using the LME method. In this case, the average coherence (1-5 Hz) between speech envelope and EMG signal was calculated for each experimental condition (loudness and syllables) and entered into a fixed-effect model with loudness, syllable, and EMG as factors.

### 2.9. Data availability

The data used in the current study will be available from the corresponding author upon request based on a formal data sharing agreement with Prof. Joachim Gross. The Matlab code of the proposed artifact rejection approach is publicly accessible through the Open Science Framework (https://osf.io/uc8st/).

## 3. Results

### 3.1. EXP1-Head movement analysis

To investigate whether speech production causes head movement during MEG recording, we extracted continuous head localization information which was captured by fiducial coils. We extracted the head center movement information for different experimental conditions, i.e. different loudness: No-sound/Normal/ Loud as well as different syllables: Pa/Du/La. Fig. 2A illustrates the Z-axis grand averaged head movement during different syllables pronunciation. These results show that, independently for each syllable, the amount of head movements is directly linked to its loudness level. Next, we calculated the averaged head movement values for the first 4 seconds after the syllables pronunciation onset. We normalized the averaged value for every participant by subtracting the No-sound condition from other experimented loudness. Boxplots in Fig. 2B display that higher loudness caused larger Z-axis head movements. We tested for the effect of loudness and syllable on head movements/ orientations by performing a linear mixed-effect model. We computed the averaged head movement/ orientation in all directions (i.e. X/Y/Z/Ox/Oy/Oz) for the first 4 seconds after the syllables pronunciation onset. All the average head movement/orientation values were entered into a fixed-effect linear model with loudness and syllable as factors. Our results show that the main effects of loudness were highly significant on the head movement in Z direction (t(98) = 5.9, p < .0001). Detailed results are presented in Table. 1.

**Table 1.**
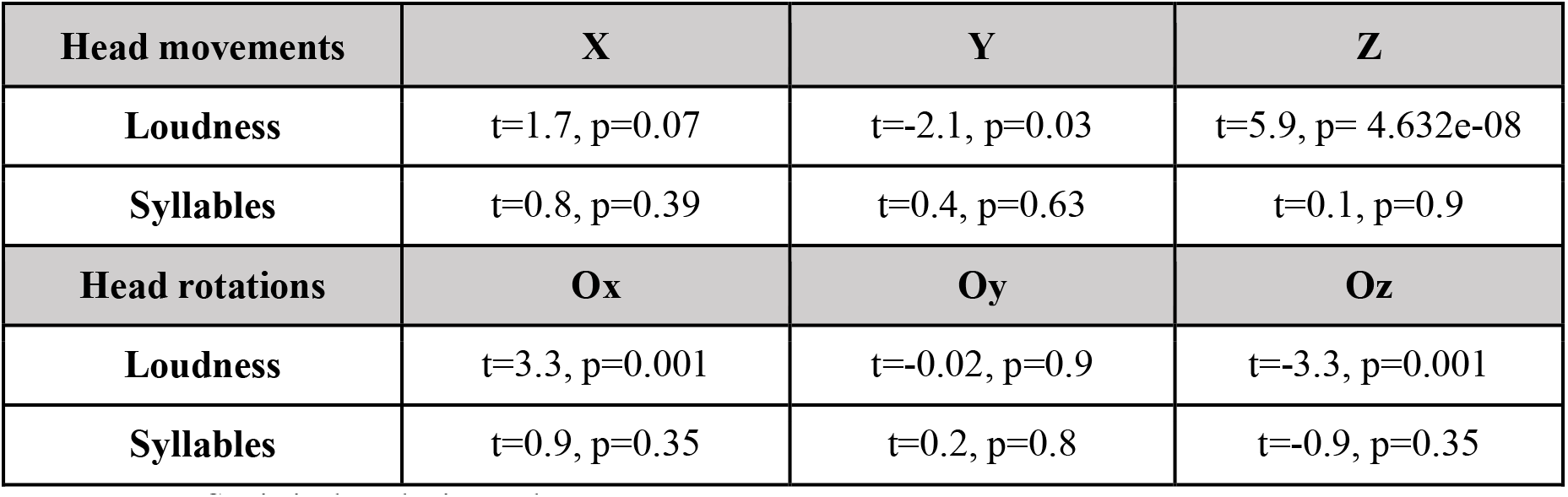
Statistical analysis results.

| Head movements | X | Y | Z |
| --- | --- | --- | --- |
| Loudness | $t=1.7$, $p=0.07$ | $t=-2.1$, $p=0.03$ | $t=5.9$, $p= 4.632e-08$ |
| Syllables | $t=0.8$, $p=0.39$ | $t=0.4$, $p=0.63$ | $t=0.1$, $p=0.9$ |
| Head rotations | Ox | Oy | Oz |
| Loudness | $t=3.3$, $p=0.001$ | $t=-0.02$, $p=0.9$ | $t=-3.3$, $p=0.001$ |
| Syllables | $t=0.9$, $p=0.35$ | $t=0.2$, $p=0.8$ | $t=-0.9$, $p=0.35$ |

**Fig. 2.**
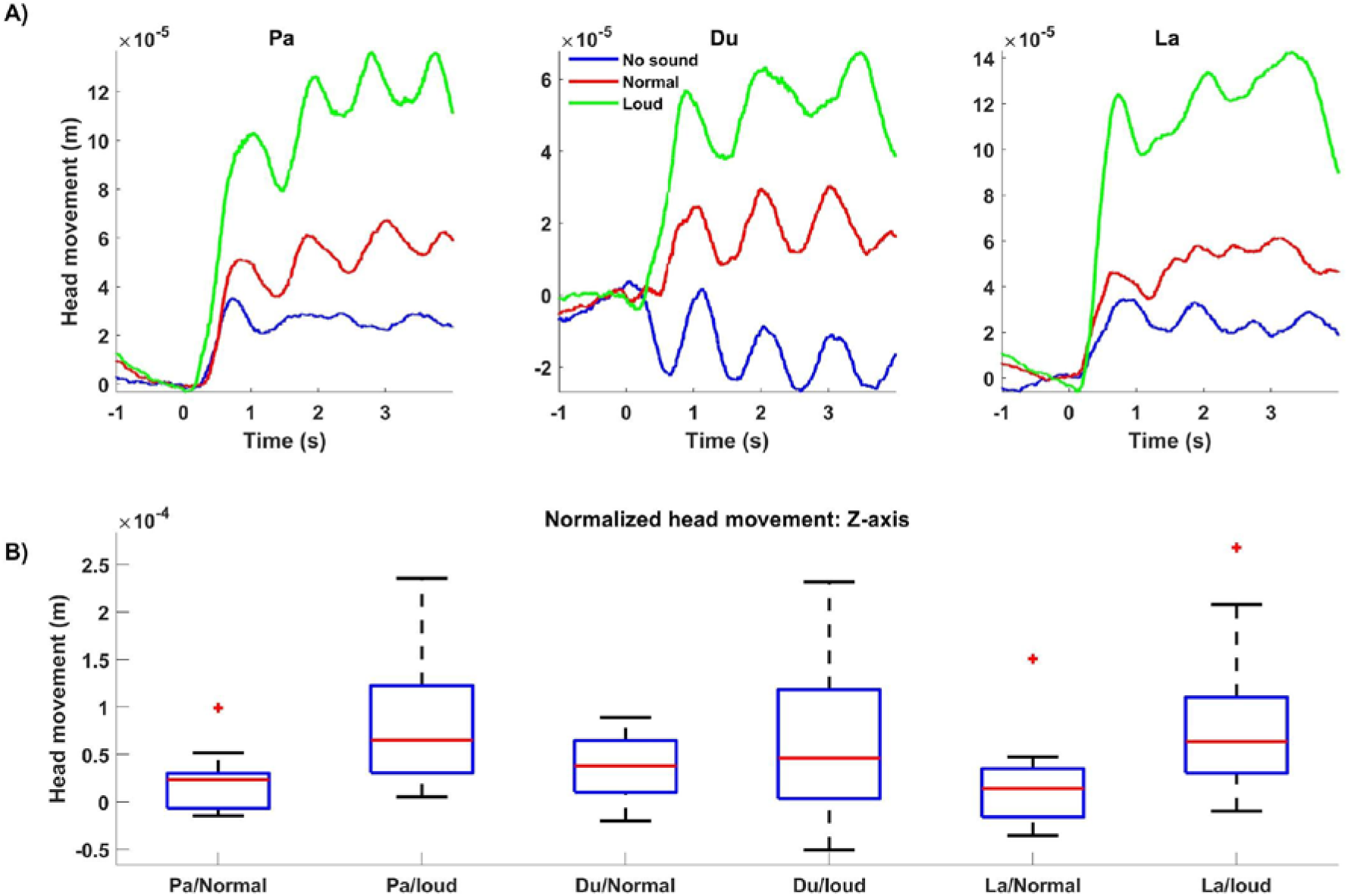
Grand averaged head movement modulated by different syllables and speech loudness. A) Head movement in the Z-axis direction is plotted for all three syllables: Pa (left panel), Du (middle panel), and La (right panel) with different loudness levels (No-sound (blue), normal (red) and loud (green)). The level of head movement is directly linked to the speech loudness. B) Boxplots showing the distribution of averaged head movement, for the first four seconds after pronunciation onset, normalized by subtracting No-sound condition from other loudnesses. For all the three syllables, higher loudness caused the stronger head movement.

### 3.2. EXP1-Speech envelope

For further analysis, we calculated the speech signal envelope. We observed a strong temporal alignment of the speech envelope with head position time-series, MEG, and EMG signals recorded from the chin, lip, and cheek during syllables pronunciations (Fig. 3A). The strong temporal alignment between these signals shows that syllable pronunciation could cause head movement and subsequently MEG distortion. In the next step, we aimed to investigate the temporal relationship between the speech envelope and EMG signals to better understand how they both reflect speech production artifacts in MEG. We therefore calculated coherence between both signals in order to identify the EMG signal showing speech-related information the best. We normalized the coherence value for every participant/syllable by subtracting the No-sound condition from other experimental conditions. Next, we averaged the normalized coherence values across syllables for every participant. We observed the strongest coherence between EMG Lip signal and speech envelope (Fig. 4A). Hence, further analysis in this study is done by using EMG Lip signals. We also expected delays between the movement of articulators and onset of speech because auditory and visual speech are temporally coupled (Chandrasekaran et al., 2009; Park et al., 2018, 2016) but typically mouth movements precede auditory speech (Schwartz et al., 2004). Therefore, we computed coherence between EMG and time-shifted speech envelope signal to identify the optimal delay. Fig. 4B illustrates the grand averaged coherence for delays between −333 and 333 ms. EMG and speech envelope showed the highest coherence at a negative delay of around −150 ms, indicating, as expected, that movement of articulators precedes auditory speech. Optimal temporal delays varied across participants (Fig. 4C). Therefore, for further analysis, the mean optimal temporal delay across the group (−150 ms) was taken into account and the coherence values were recalculated after shifting the speech envelope. The new coherence results illustrate high coherence in normal and loud speaking conditions (Fig. 4D). We tested for the effect of loudness, syllable, and EMG signal on coherence between EMG and time-shifted speech envelope signal by performing a linear mixed-effect model. We computed the average coherence in the frequency range of 0-5 Hz. All the average coherence values were entered into a mixed-effect model with loudness, syllable, and EMG signal as factors. Our results show that the main effects of loudness and syllable were highly significant (Loudness: t(290) = 9.5, p < .001; Syllable: t(290) = −4.3, p < .001; EMG: t(290) = −1.2, p = 0.2).

**Fig. 3.**
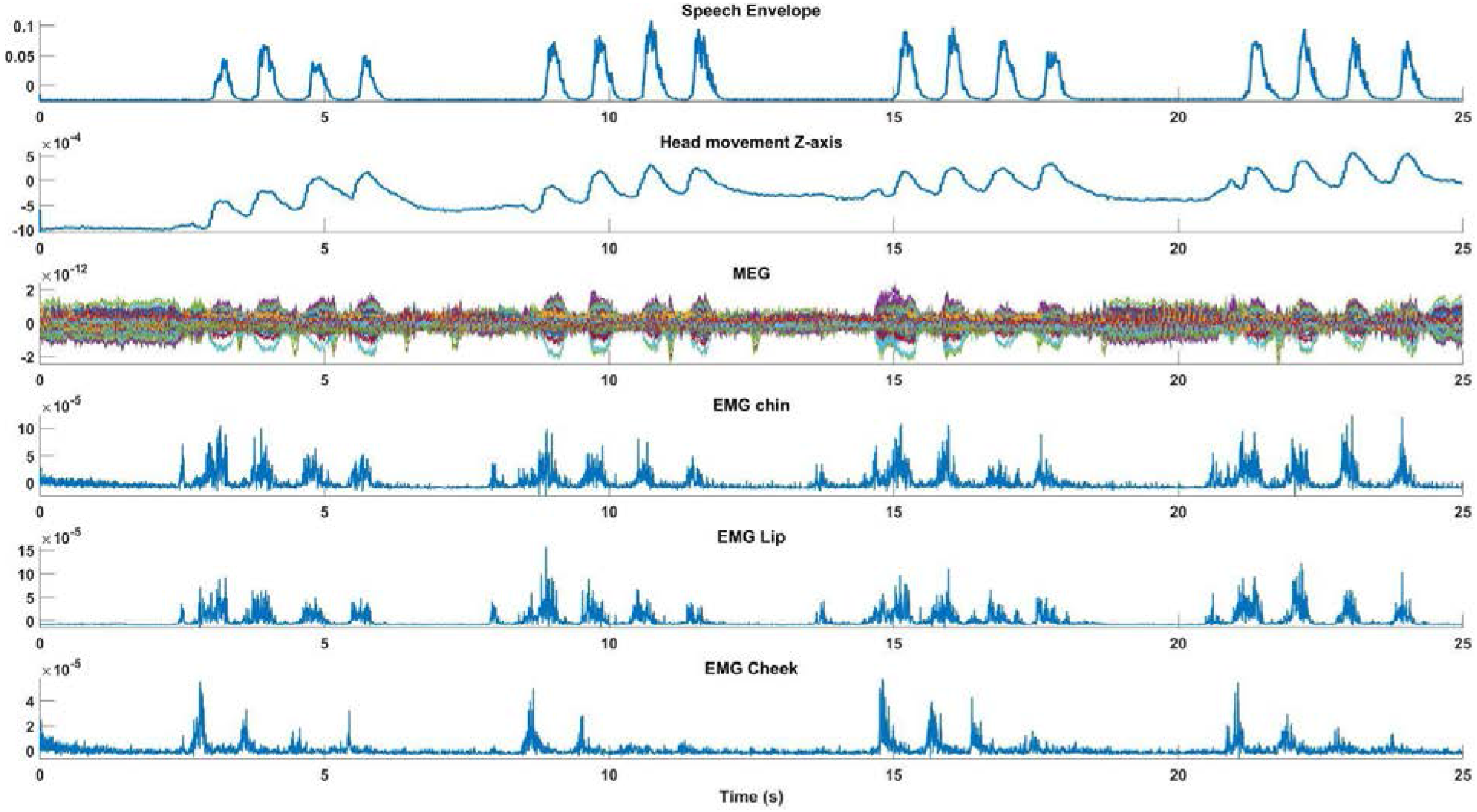
Temporal alignments between speech envelope and MEG. The first 25 seconds of an individual time-series from MEG, EMG, speech envelope, and head movements are illustrated for condition Pa/Loud. The strong temporal alignment between these signals shows that syllable pronunciation causes head movements and results in MEG distortion.

**Fig. 4.**
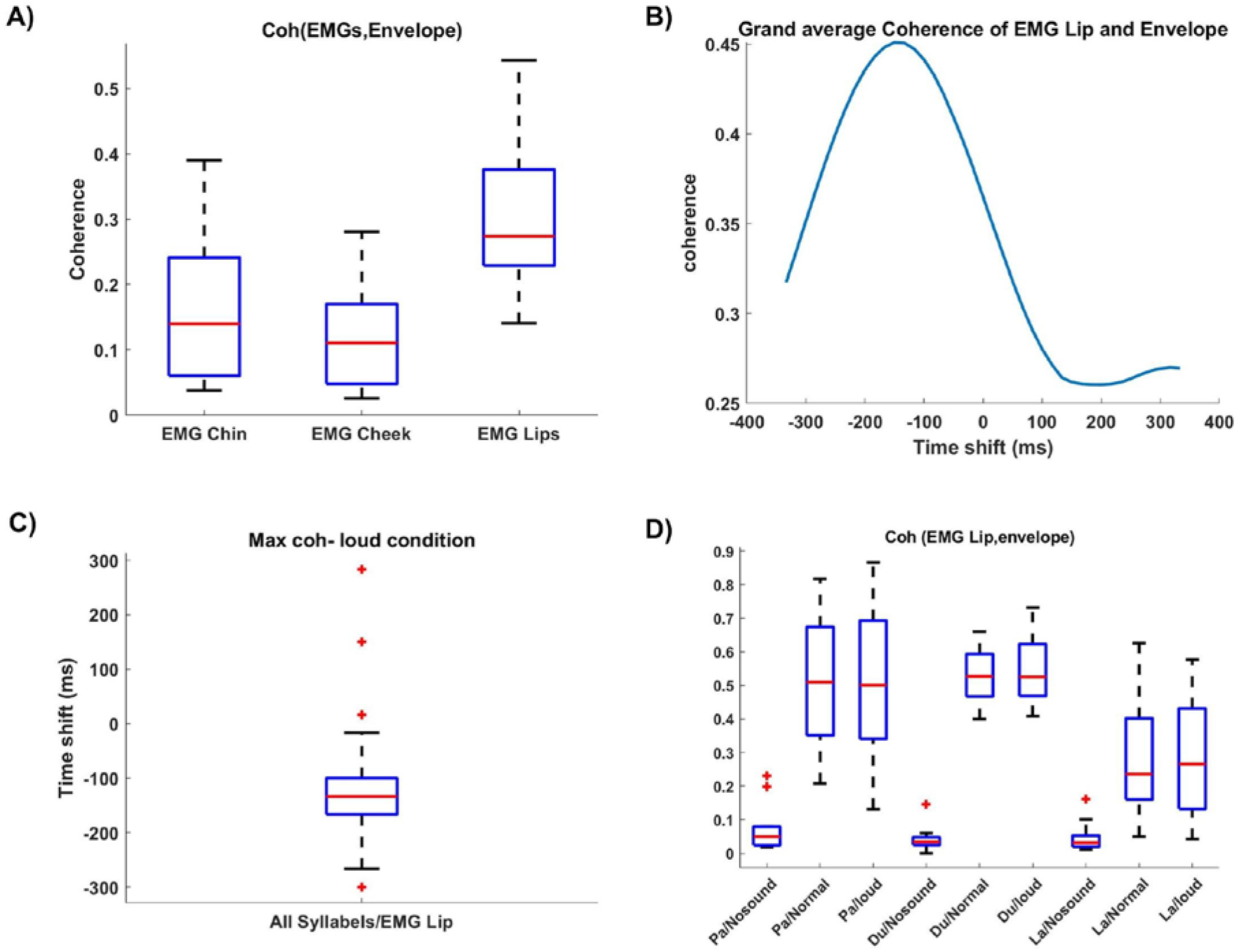
Coherence between recorded EMG and speech envelope. A) Boxplots showing the distribution of the coherence, between EMG signals and speech envelope, normalized by subtracting the No-sound condition from the loud condition and averaged across all syllables. EMG Lip signal displays the strongest coherence with the speech envelope signal. B) Coherence was computed for each participant between the EMG signals and time-shifted speech envelope. These coherence values were subsequently averaged across all participants for each pair (EMG, Envelope) and each frequency. Finally, for each pair, the coherence values were averaged within the frequency range 0-5 Hz. The group results show the optimal temporal delay in −150 ms. C) The distribution of optimal time-shifts leading to maximum coherence across participants. D) Boxplots display the distribution of coherence between the shifted speech envelope (−150 ms) and EMG Lip for different syllables and loudness. For all the three syllables, normal and loud loudness caused stronger coherence in comparison to the No-sound condition.

### 3.3. EXP1-Recording MEG data during speech production

Speech production caused head movement in all eleven participants. Subsequently, clear patterns of MEG distortion were observable for all the participants due to their head movement. We observed severe MEG distortion especially for participant #11 due to retainers (Fig. 5, first panel). Therefore, here we first report the results from this participant to study to what extent we can remove the artifact. A successful artifact correction should ideally lead to data where auditory evoked responses to speech onset can be seen. This is particularly challenging in our case where self-produced speech leads to attenuation of auditory evoked responses (Houde et al., 2002).

**Fig. 5.**
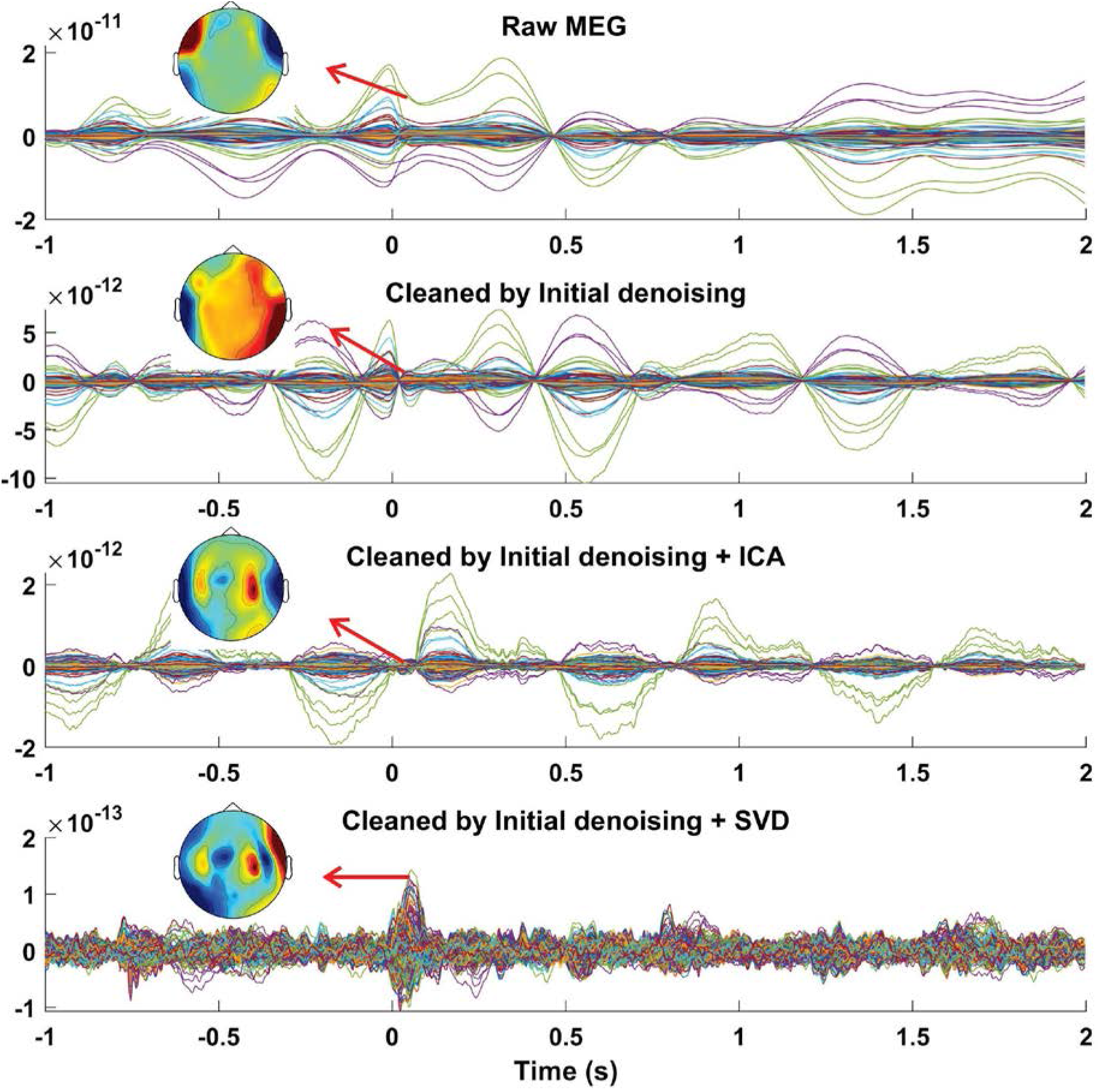
MEG recording during speech production. Averaged, time-locked data across trials of all MEG sensors from one representative individual (participant #11) recorded during syllables pronunciation before artifact rejection (first panel), after cleaning by our initial denoising step, i.e. GLM-based analysis (second panel), after cleaning by ICA (third panel), and after cleaning by SVD (fourth panel). The distribution of the averaged MEG amplitude for a time period between 50 to 60 ms is displayed on the topographical maps for different conditions. The first row illustrates strong MEG data distortion which is distributed mainly in the temporal areas. The third row shows that ICA could not successfully remove the artifact and signals of interest are still not observable due to the residual artifacts. The fourth row, however, displays a clear pattern of neurophysiological activity after removing artifacts on the topoplot. Timepoint 0 s marks syllable pronunciation onset. The data shown here are from a participant with a retainer.

Our initial artifact rejection step, the GLM-based approach, could reduce the confounding variance that was caused by head movement (Fig. 5, second panel). We tried to further remove the induced artifact from the recorded MEG data using both ICA and our proposed method. After decomposing MEG data to independent components (ICs), we observed that ICA could not successfully decompose the data due to the non-stationary nature of the induced artifact. Artifact patterns were observed in most of the components’ topographical maps and time-series. Therefore, we only removed the first four components (out of 20) containing the major part of the artifact. Although the neurophysiological pattern (auditory evoked response) was partly detectable shortly after the pronunciation onset, the residual artifact still significantly distorts the back-transformed MEG (Fig. 5, third panel). Next, we used our proposed method to remove the speech-related artifacts from the recorded MEG data. Only four components were classified as artifactual components based on our criteria. After removing artifactual SVD components, we observed a clear auditory evoked response for data recorded from participant #11 (Fig. 5, fourth panel).

We applied our proposed artifact rejection approach to clean all the recorded datasets in *EXP1* (participant #1 to #10). On average, 1.4±1.1 components were classified as artifactual (MI mean=0.16, MI SD=0.04, range=0.08-0.23) and 18.06±1.17 components (MI mean= 0.04, MI SD= 0.02, range=0.01-0.13) were classified as components containing signal of interest.

To evaluate if our artifact rejection approach could remove artifacts and retrieve neurophysiological information, we investigated if we could detect M100 activity before and after artifact rejection.

Before any artifact rejection, it was not possible to detect M100 peak due to the data distortion around the syllables production onset (Fig. 6A, left panel). However, after artifact removal, neurophysiological activities were detectable 50 ms after the syllables production onset on the grand average data (Fig. 6A, right panel). The strongest MEG distortion was observed around the syllables pronunciation onset and our artifact rejection approach was capable of largely removing the artifact (Fig. 6B and C). Residual artifact level was computed for the group data after different cleaning steps, i.e. Raw, GLM, and GLM+SVD. First, we tested for the effect of different artifact correction on the residual artifact level by performing a linear mixed-effect model. We computed the level of the artifact by applying the constructed spatial filter on data. All the artifact level values were entered into a fixed-effect model with method as factor. Our results show that the main effects of artifact corrections were highly significant (Methods: t(28) = −4.4, p =0.0001). Our results showed that our proposed approach could significantly remove the artifact from the recorded data (Fig. 6C, right panel).

**Fig. 6.**
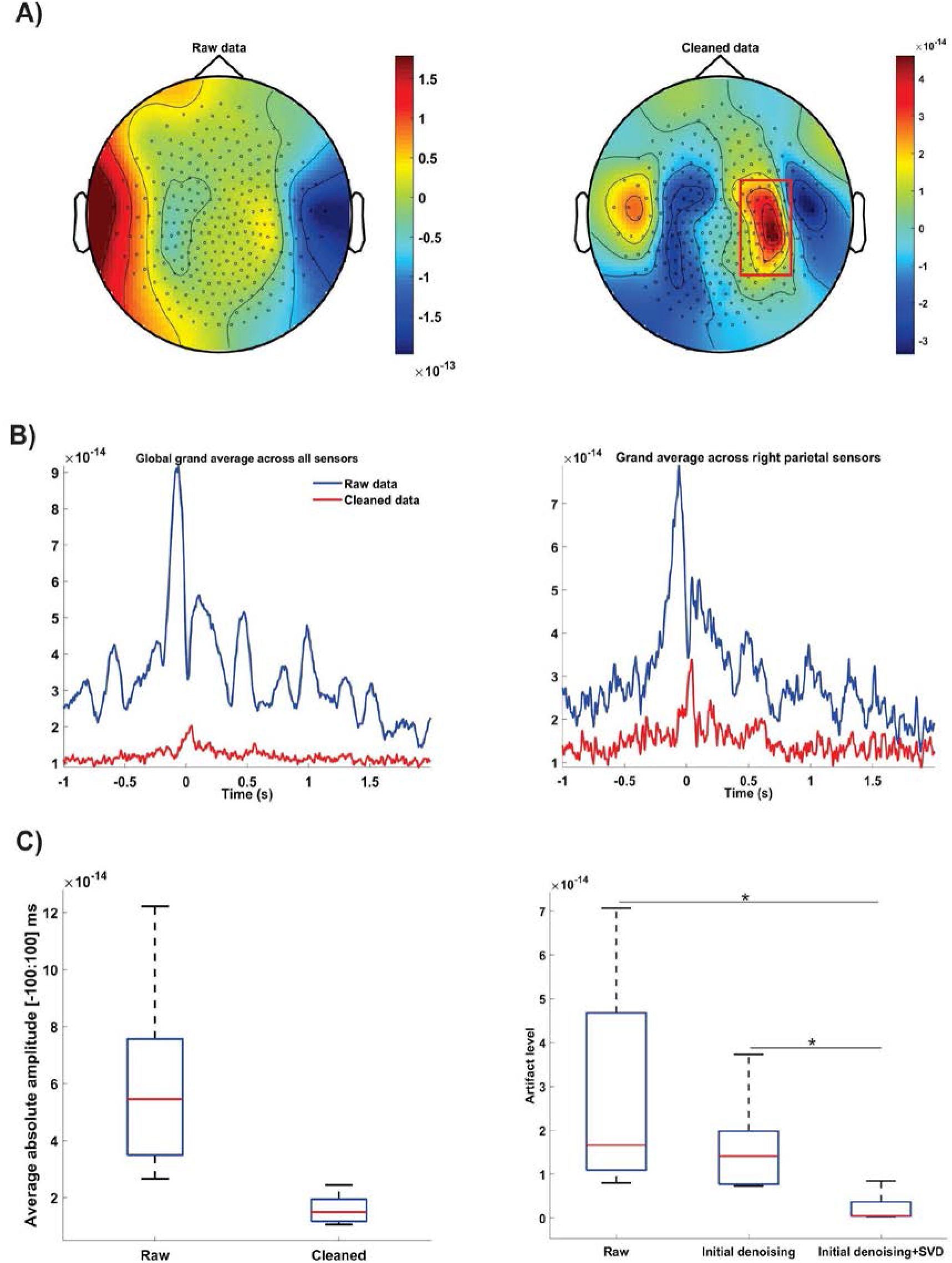
MEG artifact rejection. A) Average topographic distribution of MEG amplitude between 40 and 50 ms for raw data (left column) and after cleaning data by our proposed method (right column). M100 activity was masked before artifact rejection by speech related artifact. Color codes MEG amplitudes. B) Absolute grand average of the time-series from all MEG sensors (left column) and sensors above the right parietal area (right column) from all the participants before (blue) and after (red) artifact rejection. Selected sensors are designated by a red rectangular C) Boxplots (left panel) showing the distribution of the average values of MEG amplitude between −100 and 100 ms across participants for raw and cleaned data. Right panel boxplots illustrate the distribution of artifact level after data cleaning by different approaches. Our proposed method could significantly remove artifacts from MEG data in comparison to other methods. Significant P values (after t-test analysis) are indicated. * P < 0.01.

### 3.4. EXP2-Recording MEG data during syllable perception

We also evaluated if the proposed artifact rejection approach partly removes neurophysiological information. We applied the spatial filters constructed from the speech production dataset to the syllables perception dataset in *EXP2.* Since there is no speech artifact in the perception condition an ideal artifact correction would leave the perception data unchanged. We observed severe MEG distortion for participant #9 due to retainers in the speech production condition. Six components were classified as artifactual for this dataset. We constructed a spatial filter from these six components and applied it to the syllables perception condition. We observed a clear M100 peak in response to syllable onset before (Fig. 7A) and after (Fig. 7B) artifact correction steps. Importantly, the auditory topography was not attenuated or distorted after artifact correction. Instead, small artifacts in temporal and frontal areas were reduced leading to an overall cleaner topography. Next, we quantified the performance of our artifact correction at the group level. We constructed the SSP filters for every participant in *EXP2* from the speech production dataset. After applying the constructed filter to syllable perception dataset, we observed that the proposed method leads to negligible changes of the M100 amplitude (median: 0%; 25th percentile: −5%; 75th percentile: 0.7%; Fig. 7C and D).

**Fig. 7.**
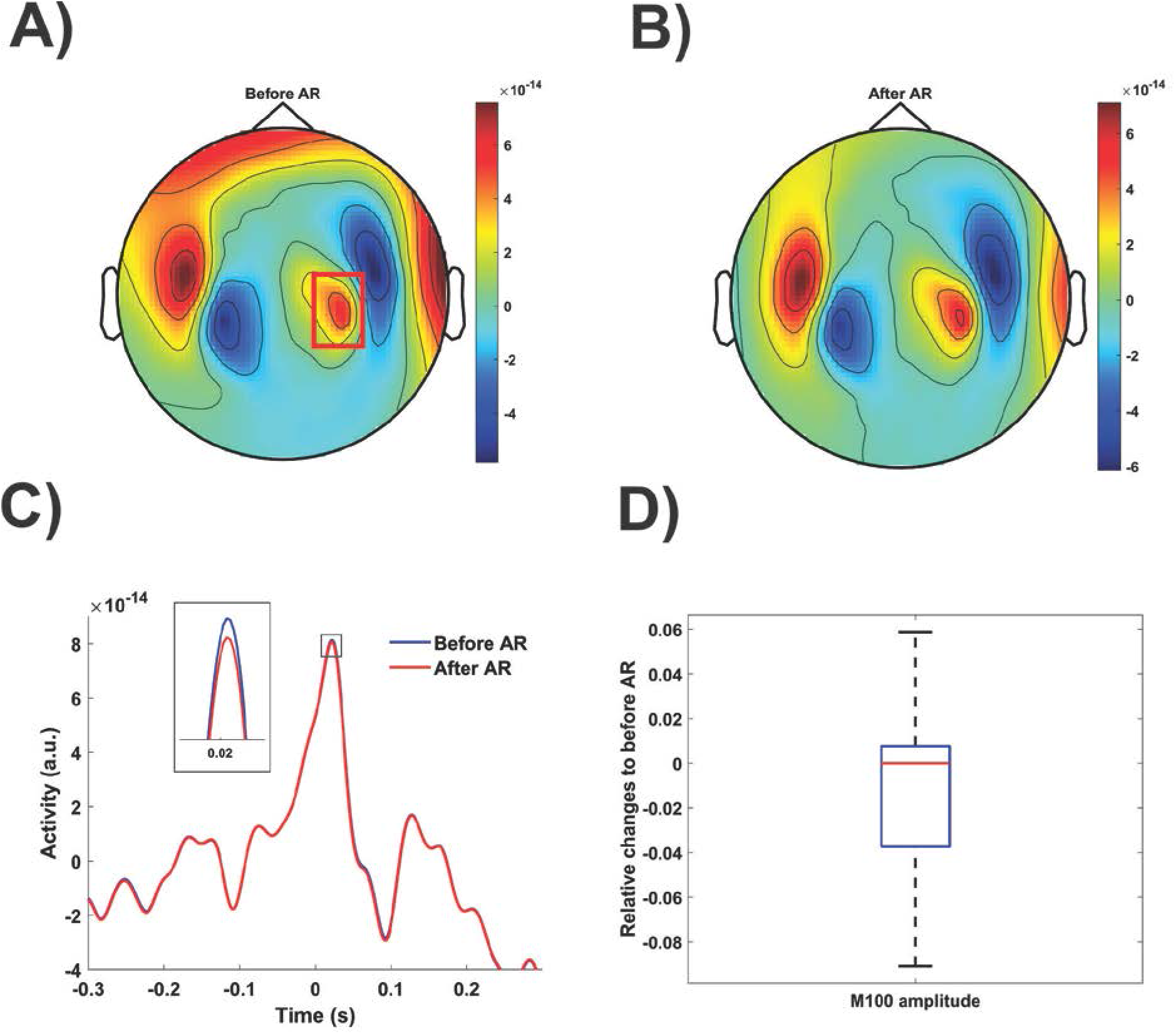
MEG artifact rejection on syllables perception condition. Topographic distribution of MEG amplitude between −20 and 70 ms for raw data (A) and after cleaning data using spatial filters constructed from speech production dataset (B) for participant #9 of *EXP2.* Color codes MEG amplitudes. C) Grand average of the time-series from the MEG sensors, designated by a red rectangular, from participant #9 before (blue) and after (red) artifact rejection. The proposed artifact rejection caused slight M100 amplitude removal. D) Boxplots showing the distribution of the relative change of individual M100 amplitudes after artifact rejection relative to before artifact rejection condition for all the participants in *EXP2* (AR=artifact rejection).

In summary, this indicates that the SSP correction specifically targets the speech artifact and leaves the real neurophysiological signals of interest largely intact.

## 4. Discussions

In this study, we carefully characterize the effects of head movement on the recorded MEG data during speech production. We demonstrate that MEG data is distorted due to head movement and its severity is directly linked to the speech loudness level. We present an artifact rejection approach to remove artifacts caused by head movement from MEG data. Our approach is based on a combination of regression analysis and signal space projection (SSP). The proposed method was successfully applied to the MEG data recorded during pronouncing syllables with different loudness.

Using the real-time head center positions extracted from the head localizers coils, we were able to accurately track head positions during different experimental conditions. We observed that the head movement level is directly linked to the loudness level of the pronunciation of the syllables. This loudness-level-dependent movement can easily distort the recorded MEG data during speech production studies. In addition to general head movement, the unique pattern of lower jaw movement and facial articulatory muscles, for producing every single word, add a complex artifact pattern to the recorded data especially when participants have some ferromagnetic implants in their mouth such as retainers. Moreover, we observed a strong connection between speech envelopes and the facial articulatory muscles involved in speech production. Taken together, there are several different sources that contribute to MEG distortion during speech production studies. One of the major sources is the head and lower jaw movement. In this study, we mainly focus on removing this type of artifact.

A few earlier studies have investigated speech production using MEG. They used different strategies to either avoid or remove the speech-related artifacts. In the following, we discuss these approaches and compare them with our proposed artifact rejection approach.

Earlier studies have avoided recording neurophysiological activities during speech production due to data distortion. Some tried to simply circumvent the induced artifacts by delaying continuous speech, silent naming, or manual responses (Eulitz et al., 2000; Ewald et al., 2012; Liljeström et al., 2008; Sahin et al., 2009; Schmitt et al., 2000). However, these approaches could prevent us from directly investigating neural activities underlying speech production. In another study, (Saarinen et al., 2006) reported the feasibility of investigating cortical oscillations around 20 Hz using MEG without artifact rejection. While they reported reliable observations, this approach might be only feasible for special cases such as extracting some features which are not affected by speech-related artifacts. Using EEG, other studies tried to investigate speech-related brain activity by removing artifactual epochs (Ewald et al., 2012) or by using heavy low-pass filtering (Masaki et al., 2001). While these approaches are transferable to MEG, they are not applicable when we record data during continuous speech tasks, which could be distorted by head movement continuously, especially in lower frequency bands.

Temporal signal space separation (tSSS) (Taulu and Simola, 2006) is an MEG artifact rejection approach that has been used in several studies to reduce speech artifacts in MEG data. The basic idea behind this approach is to separate MEG signal components originating inside the sensor arrays from sources outside the brain using the sensor geometry and expansions of spherical harmonic functions. Several previous speech studies have shown that after correcting head movement and suppressing external disturbances by tSSS they could investigate corticomuscular coherence, cortical neural network connectivities as well as the connection between evoked responses and rhythmic activities during speech production (Laaksonen et al., 2012; Liljeström et al., 2015b, 2015a; Ruspantini et al., 2012). However, since most high amplitude speech-related artifacts come from sources such as facial muscle activity or lower jaw movement (which are inside the sensor array), tSSS could not completely clean the data. Therefore, several complementary strategies have been taken to further clean the data after applying tSSS. Ruspantini and colleagues trained their participants in order to produce small movements during speech production (Ruspantini et al., 2012). Liljeström and colleagues tried to avoid the residual artifact by selecting artifact-free time windows for studying cortical networks underlying preparation for speech production (Liljeström et al., 2015a). Finally, in recent studies, the combination of tSSS and independent component analysis (ICA) has been used to remove speech-related artifacts from the recorded MEG data (Alexandrou et al., 2017; Bourguignon et al., 2020). In ICA, the spatial filters are derived by decomposing sensor data to the set of maximally temporally independent components (Delorme et al., 2007). Alexandrou and colleagues used ICA to remove artifacts induced by the movement of facial articulatory muscles during speech production. Their results showed that they observed artifacts separated into several components (Alexandrou et al., 2017). This is in line with our observation when we tried to clean data by means of ICA. For participant #11, for instance, topographical maps for all of the components showed a noisy pattern. There are several reasons why ICA might not be optimal for speech-related artifact rejection. The movement artifact induced in MEG signals is non-stationary due to the different articulation depending on the word to articulate and the location of the source of the artifact (e.g. jaw) changes over time (De Vos et al., 2010). This violates the requirements for ICA and leads to a failure of ICA to clearly separate signals and artifacts.

Taken together, different strategies/approaches have been used to study speech production using MEG. However, there is still a demand to further improve speech-related artifact removal approaches. The unique characteristic of our proposed approach is that it makes use of head movement information to initially reduce movement artifacts and then uses the signal space projection approach (SSP) to remove the residual artifacts.

In the first step of our proposed artifact rejection pipeline, we used regression analysis to reduce the artifact level. We could successfully reduce the confounding variance caused by head movement by incorporation of the head position/orientation time-series into a general linear model. This approach has been previously applied on several MEG datasets recorded during visual attention, auditory expectation, as well as in somatosensory spatial attention tasks (Stolk et al., 2013). This step could play a vital role in improving the signal-to-noise ratio by removing a major part of the artifact. It should be noted that this step benefits from running twice. First, global head movements are corrected that occur across the duration of the measurement. Second, a third-order polynomial fit to the head movement traces is subtracted to optimise the representation of transient movements (such as movements associated with each syllable production). For two out of eleven participants this step was sufficient to clean data and proceed with further analysis. The effectiveness of this step depends on the distortion level (level of confounding variance) and the quality of the extracted head position/orientation time-series.

To remove the residual artifact, we used the signal space projection (SSP) approach in the second step. Using SSP, we can separate signals into a signal subspace and an artifact subspace (Uusitalo and Ilmoniemi, 1997). Data is then linearly projected onto the signal subspace. This is an effective approach for separating artifacts from signals especially when the artifact has a constant spatial pattern. This approach has been previously used to remove different types of artifacts such as ocular artifacts from MEG data (Chang et al., 2018; Pulvermüller and Shtyrov, 2009). During speech production, movement artifacts are represented in a distinct subspace (coming from either head movement or lower jaw movement) with varying amplitudes as a function of time. Therefore, SSP is an optimal approach for separating spatial subspaces containing movement artifacts. We used SVD to decompose MEG data into components. Movement-related components were selected by a conservative three-steps approach using components’ topographical maps and time-series as well as their mutual information with head position/orientation time-series. The advantage of the latter component selection step is that it makes explicit use of the available information on artifact characteristics.

In this study, we demonstrated that our proposed approach was able to remove artifacts and retrieve neurophysiological information. Importantly, our approach could successfully clean MEG data strongly distorted due to implanted ferromagnetic dental retainers. Previous studies reported successful MEG measurements in the presence of implanted ferromagnetic stimulation hardware (Abbasi et al., 2016). Although the pre-measurement demagnetization step might reduce the artifact caused by retainers, any head/lower jaw movement can still induce strong artifacts in MEG sensors during measurement. Here we also showed the feasibility of recording participants with retainers. This could be very important for recruiting participants not only for studying speech production but also when planning any MEG experiment.

Another advantage of the proposed approach is that there is no need to perform any initial step such as defining epochs or frequency bands representing either artifact or neurophysiological signal. Moreover, this approach is system-independent and could be applied to any type of MEG data.

Additionally, facial muscle activities have been reported as another source of artifact in MEG speech studies. The power of induced muscle artifact lies in frequencies higher than 20 Hz (Muthukumaraswamy, 2013). In this study, we did not attempt to remove muscle artifact from MEG data. Previous studies used different strategies such as the ICA approach to remove muscle artifact (McMenamin et al., 2011; Muthukumaraswamy, 2013; Shackman et al., 2009). Generally, ICA shows better performance when data quality is higher. Therefore, in case muscle artifact removal is necessary, we recommend to improve the signal-to-noise ratio by means of our proposed approach and apply ICA as the final step.

## 5. Conclusion

Our newly presented artifact rejection approach could facilitate assessing speech production and perception mechanisms by MEG. The presented results revealed that it is also possible to remove complex patterns of artifact induced by implanted devices, while preserving essential neurophysiological information. Future applications might include MEG experiments during continuous speech production assessing the neural activity underlying speech production.

## CRediT authorship contribution statement

### Omid Abbasi

Conceptualization, Formal analysis, Investigation, Methodology, Software, Visualization, Writing – original draft.

### Nadine Steingräber

Data collection, Formal analysis, Data curation, Writing – review & editing.

### Joachim Gross

Conceptualization, Funding acquisition, Methodology, Project administration, Supervision, Writing – review & editing.

## Acknowledgments

We acknowledge support by the Interdisciplinary Center for Clinical Research (IZKF) of the medical faculty of Münster (grant number Gro3/001/19) and the DFG (GR 2024/5-1).

